# Modelling the spatiotemporal dynamics of multispecies population interactions in the context of climate change

**DOI:** 10.1101/2024.02.07.579316

**Authors:** Katerina Christou, Reto Schmucki, Hélène Audusseau

## Abstract

1. Species interactions are fundamental to the stability and productivity of ecosystems. To improve our capacity to predict and understand how climate change shapes the distribution of species and the dynamics of biotic interactions, we need to develop spatially explicit multi-species models that are built upon species-specific responses to changing conditions.
2. We developed a two-dimensional diffusion-advection-reaction predator-prey model that integrates species-specific responses to heterogeneous landscapes, affecting species dispersal, reproduction and survival rates. We derived conditions for the stability and feasibility of the coexistence steady-state and observed how temperature variation stabilises or destabilises the system.
3. We conducted numerical simulations to explore the effect of predicted extreme temperatures on the spatial dynamics of a parasitoid-butterfly system and their interactions. Applied to four different climatic environments, the numerical approximations demonstrate the asymmetric impact of a warming climate on interacting species. The output density distribution maps highlight the capability of our model to produce interpretable multiscale predictions which can be used to identify and evaluate species vulnerability locally and across their range.
4. By building upon a solid mechanistic understanding of species-specific responses to environmental change, our model can be extended to other species and variables, including environments where the availability of empirical data is limited, and explore the dynamics and distribution of interacting species under different scenarios of environmental change.

## Introduction

Species interactions are fundamental to the productivity and stability of ecosystems (Traill et al., 2010), supporting processes that regulate the dynamics and distribution of biodiversity (Wisz et al., 2013). Despite their ecological importance, our understanding of the impact of climate change on such interactions and how they influence population dynamics and community assemblages remains scarce (Blois et al., 2013; Wisz et al., 2013). While temperature influences all organisms, the way and extent to which species respond to change and variability in their thermal environment vary across taxonomic groups and trophic levels (Boggs, 2016; Lenoir et al., 2020; Parmesan, 2007; Poloczanska et al., 2013). Such variations in species sensitivity and response to climate change will not only affect individual species’ demography but also have consequences on species interactions and their distribution (Audusseau et al., 2017; Laws, 2017). Integrating species-specific responses to temperature in spatially explicit species interaction models will significantly improve our capacity to predict the dynamics of species and biological communities in changing environments.

To predict the impact of climate change on species interactions and biodiversity, models must focus on integrating processes that influence their capacity to adapt and thrive under new environmental conditions and, if required, to disperse to locations where conditions are suitable, e.g., higher altitude or latitude (Bellard et al., 2012). One way species respond to changing environments is through phenotypic plasticity, which enables an organism to change its behavioural, physiological and morphological traits to cope with environmental variability and enhances its growth, survival, or reproduction under specific conditions (Fusco & Minelli, 2010). The nature and extent of such responses vary greatly across species (Figure 1 in Berg et al., 2010) and are good indicators of species’ capacity to adapt to environmental changes. The difference in species’ sensitivity to temperature and capacity to adapt to changing thermal environments will also influence the strength of species interactions and their robustness to climate change (Pounds et al., 2006). When species cannot adapt to changing conditions, their survival will depend on their capacity to escape adverse environments (Watkinson & Gill, 2002) and disperse into areas where conditions are suitable (Bellard et al., 2012). Nevertheless, differences in species’ dispersal capacity (Figure 2 in Berg et al., 2010) can also result in spatial mismatches between interacting species and have direct consequences on species interactions and local survival rates (Parmesan, 2007; Visser & Holleman, 2001).

**Figure 1.**
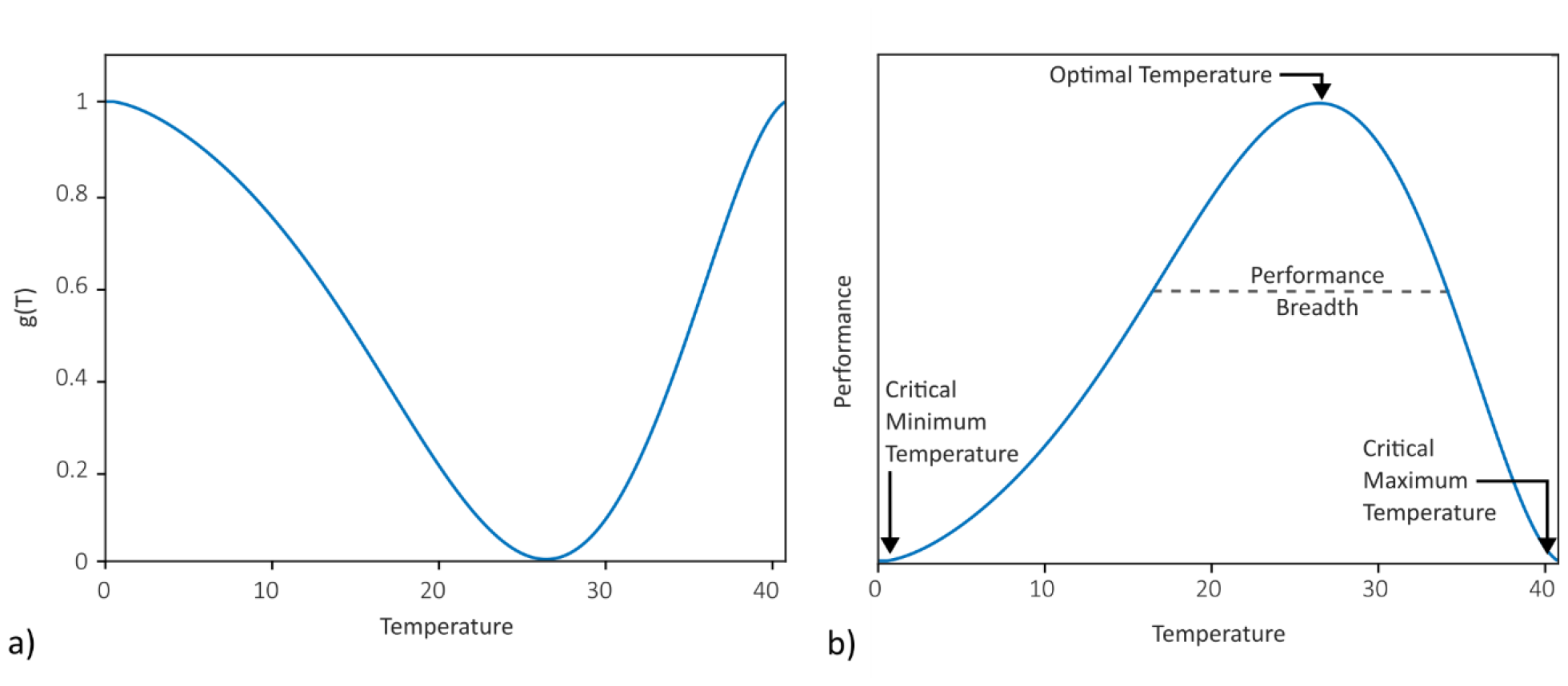
The formation of a) the climate function g(T) based on the general shape of b) the thermal performance curve for ectotherms.

**Figure 2.**
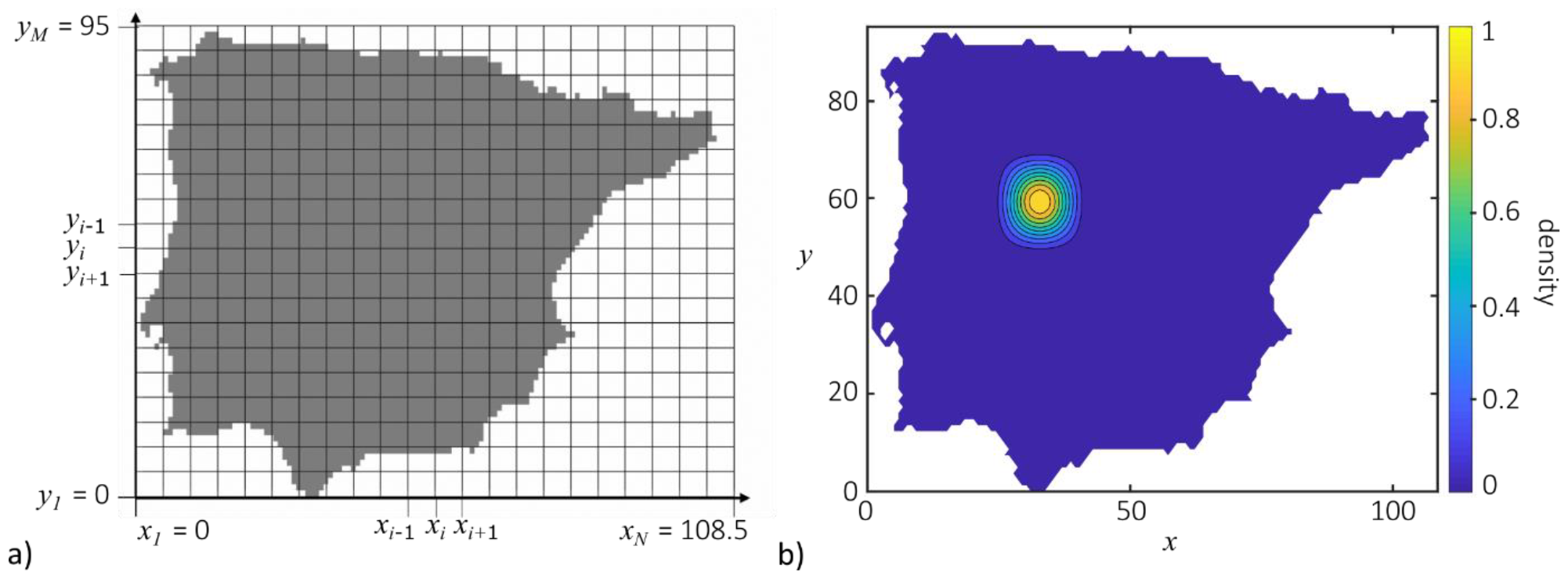
a) Two-dimensional finite difference grid over the Iberian Peninsula, where N = 130 and M = 78. The grey colour depicts the land and the white colour the sea. b) Initial conditions for the dimensionless parasitoid-host temperature-dependent system. The same initial conditions were used for both u and v, u(x, 0)=v(x, 0). The colour scale in b) shows the dimensionless population densities.

Population dynamics models provide a rigorous basis for studying species interactions and exploring different scenarios of environmental changes. Models based on ordinary differential equations have been used to explore the effects of warming on the nature and strength of the interactions between species (DeLong & Lyon, 2020; Gretchko et al., 2018; Rall et al., 2010; Sekerci, 2020; Sekerci & Petrovskii, 2015). Despite the significant contribution of these theoretical models, they do not account for the spatial aspect of species response to climate change and overlook the influence of climate on species’ dispersal, a critical factor that will shape species co-occurrence patterns.

Since the first species interaction system was formalized, the predator-prey systems based on the ordinary differential equations described by Lotka-Volterra (1920; 1926), extensions to partial differential equations (PDE) have made it possible to significantly improve the biological relevance of these models to describe not only the temporal dynamics of species interactions but also consider spatial processes. Reaction-diffusion models, initially proposed by Turing (1952) and introduced in ecology by Segel & Jackson (1972), allow the species to be modelled as diffusive. Particularly the diffusion term describes the movement of species from a region of high density to a region of lower density in a homogeneous environment. However, reaction-diffusion models, where dispersal relies solely on random diffusion, fall short of capturing the complex dynamics of species in heterogeneous environments (Rowell, 2009). Following studies, mostly theoretical, have accounted for non-random diffusion through various forms of advection terms known as cross-diffusion, self-diffusion, prey-taxis and fitness-dependent dispersal (Cantrell et al., 2013; Cosner & Winkler, 2014; Lou & Ni, 1996; Shigesada et al., 1979). For example, Grindrod (1988, 1991) addresses this limitation of reaction-diffusion models by adding a bias velocity term. This term, which describes the directed movement of species, is modelled separately from random diffusion by an advection term, in which species move along a fecundity gradient. The resulting diffusion-advection-reaction model has the capacity to integrate realistic species’ movement in a spatially heterogeneous landscape, accounting for variation and spatial patterns in resource availability and species interactions. The dynamics of the model with directed motion were considered in one-dimensional spatial domains in Grindrod (1988, 1991), and extensions to two-dimensional spatial domains were studied first for a system of two interacting species (Kurowski et al., 2017) and then for multispecies systems (Taylor et al., 2020). These developments on non-random dispersal applied to two-dimensional heterogeneous environments open new opportunities for ecologists, theoretical and empirical, to develop and apply more realistic mechanistic mathematical models to study the impact of environmental change on species interactions and investigate the urgent question of the impact of climate change on the dynamics and distribution of species. Despite the relevance of reaction-diffusion-advection models to the study of species interactions in heterogeneous landscapes, such models have not yet been applied to study and predict the effects of climate change on species dynamics. This is partly due to the challenge of integrating species-specific responses (e.g., thermal tolerance) into mathematical models. Although PDE models are fundamental to the development of realistic mechanistic models for predicting population dynamics, species interactions and many other ecological processes in space, the complexity associated with the numerical approximation of these intricate systems, particularly within two-dimensional spatial domains, requires knowledge of numerical methods that are often an obstacle for many ecologists.

Here we extend existing models to a two-dimensional diffusion-advection-reaction model for predator-prey systems that integrates species-specific responses to local climatic conditions. We account for species’ directed motion through a temperature and fitness-dependent dispersal, similar to the one presented in Grindrod (1991) for competitive species, and we also account for the effect of local climatic conditions on the carrying capacity and reproduction rate of the prey. Climate impact is introduced through a climate function based on the thermal performance curve of ectotherms. We also go beyond the existing theoretical studies by numerically solving this reaction-diffusion-advection predator-prey system in two dimensions and applying our solution to real landscapes. Specifically, we explore the impact of past and future climates on a host-parasitoid system (butterflies and wasps) in the Iberian Peninsula (Spain and Portugal). The numerical approximations for this system illustrate the relevance of our model to predict and explore how climate change and extreme temperature can impact insect demography and their trophic interactions. Extension of this spatially explicit diffusion-advection-reaction model is straightforward and can be applied to model the dynamics of species interactions in a wide range of environmental contexts and biological systems.

## Method

### Two-Dimensional Predator-Prey model

We propose a diffusion-advection-reaction model that allows the modification of species’ dispersal in response to their interactions and the condition of their physical habitat.

The population density of the prey species, denoted by *u*, is modelled as having logistic density-dependent growth, with intrinsic growth rate *r* and carrying capacity *K*. The population density of the predator is defined by *v* and the predation is modelled by the Holling Type II functional response, by which the rate of prey consumption by a predator rises as prey density increases, but eventually levels off at an asymptote at which the rate of consumption remains almost constant regardless of increases in prey density. (Hassell et al., 1977; Holling, 1959). Type II functional responses are the most frequently studied and well-documented in empirical studies (Dawes & Souza, 2013; Gentleman et al., 2003; Jeschke et al., 2002; Skalski & Gilliam, 2001). The rate of predation is denoted by *α* while *A* is the prey population size at which the growth rate of the predator is half its maximum. In our parasitoid-host system (butterflies and wasps), the constant *ε* represents the number of viable eggs that a parasitoid (predator) lays on a single host (prey). In general terms, *ε*is a dimensionless parameter that describes the conversion efficiencies of preys (hosts) to predators (parasitoids) and is given as a fraction or decimal. The dispersal coefficients *δ*_*u*_ and *δ*_*v*_ of the two species are constants and determine the rate at which each species disperses randomly through the domain. Finally, predators are subject to intrinsic mortality rates *d*.

The main difference compared to standard population models is that the random motion of individuals is biased by an optimal velocity *v* (*v*_*u*_ for the prey and *v*_*v*_for the predator). This velocity is selected to increase individual’s net reproduction rate. On average, the species disperse in the ideal direction. Hence the average velocity of the individuals of each species, *w*_*ρ*=*u,v*_, is the sum of their optimal velocity and the diffusive term,

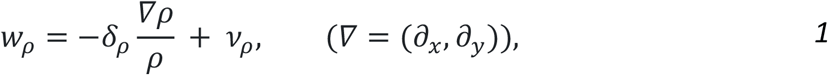

where *ρ*=*u, v*.

Species net rate of reproduction *E*_*ρ*=*u,v*_, is a constructed term comparable to the reaction terms in each species corresponding equation. A relationship between *E*_*ρ*_ and *v*_*ρ*_ is constructed with *v*_*ρ*_ as a local average of ∇*E*_*ρ*_. For convenience, we use the substitution ∇*ϕ*_*ρ*_=*v*_*ρ*_. The chosen forms have the advantage of allowing us to impose zero flux boundary conditions on *v*_*ρ*_ to prevent individuals from moving across the boundary of the domain.

As in Grindrod (1991), we define the relationship between *E*_*ρ*_ and *ϕ*_*ρ*_ of each species to be

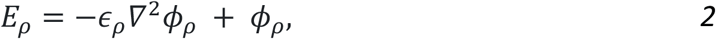

where *ρ*=*u*, v. The parameter *∈*_*ρ*_ is a very small parameter for smoothing any sharp variations in *E*_*ρ*_ and *ϕ*_*ρ*_. Conceptually, *ϕ*_*ρ*_ represents the attractiveness of a location for an individual, considering both survival chances at that specific location and the conditions in the surrounding area.

The full reaction-diffusion-advection system is thus as follows:

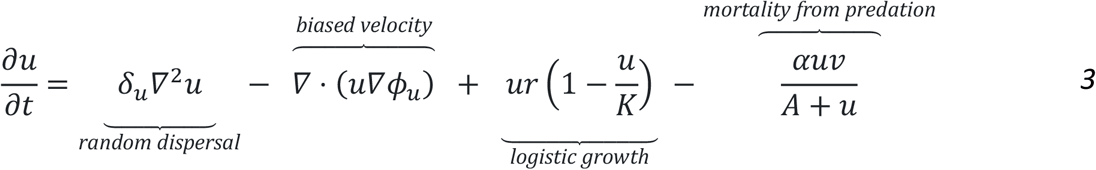

and

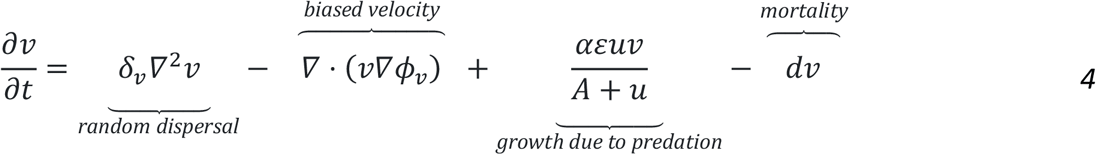

where the expressions for *E*_*u*_(*u, v*) and *E*_*v*_(*u*) are

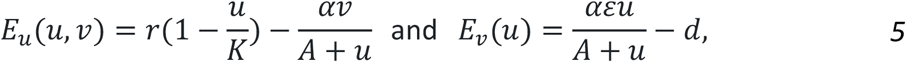

respectively.

Then, the expression for the velocity potential *ϕ*_*ρ*=*u,v*_ is

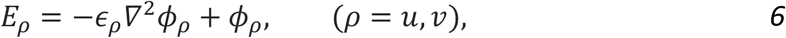

where 0 ≤ *∈*_*ρ*_ ≪ 1.

The system is posed on a given domain say Ω of lengths *L* and *P*, i.e., Ω=(0, *L*) × (0, *P*), with zero-flux Neumann boundary conditions on the boundaries *∂*Ω to close the system.

### Climate Function

We extend the work of Gretchko et al. (2018) and implemented climate functions in the two-dimensional diffusion-advection-reaction predator-prey model described above. With this climate function, we can explicitly account for the effect of temperature variation on predator-prey dynamics. The climate function describes the relationship between species’ relative fitness and temperature. This function mirrors the bell-shaped curve associated with the general thermal performance curve of ectotherms, with optimum and lower or upper critical temperatures (*Figure 1*). The inclusion of the climate function *g*(*T*) takes values from 0 and 1, where the value 0 corresponds to the most favourable temperature and 1 the least favourable (*Figure 1*).

The climate function is set to affect the velocity of both species while only affecting the prey logistic growth directly. Following *Equation 1*, depicting the velocity by which the two species populations are dispersing, the temperature-dependent velocities of the prey and the predator are now given by

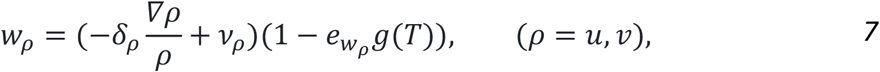

where 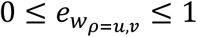 are scaling factors of the climate function.

Similarly, the intrinsic growth rate and the carrying capacity of the prey are dependent on the temperature by the equations

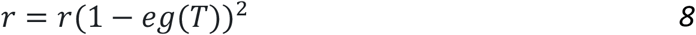

and

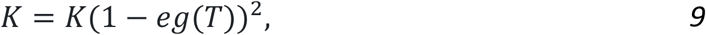

respectively. Again, 0 ≤ *e* ≤ 1 *i*s a scaling factor of the climate function.

### Temperature-dependent two-dimensional predator-prey model

The climate function defined above (*Equations 7-9*) is added to the full reaction-diffusion-advection system (*Equations 3 and 4*) and the two-dimensional predator-prey model with temperature dependence is given by

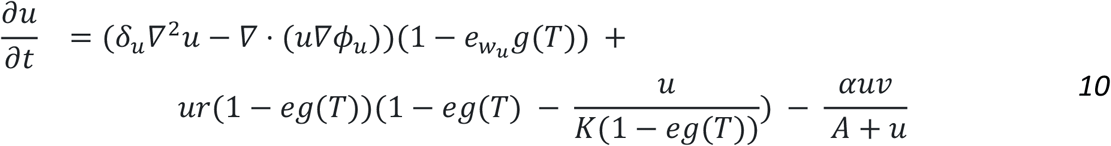

and

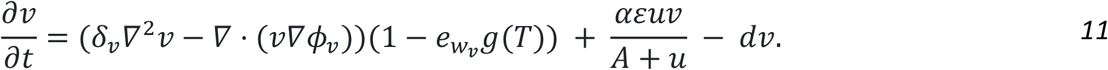

The expressions for *E*_*u*_(*u, v*) and *E*_*v*_(*u*) are given by the temperature-dependent system (*Equations 10 and 11*) in the form

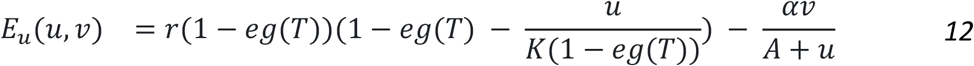

and

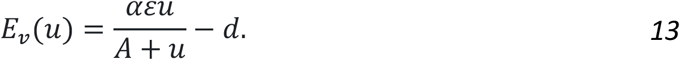

As already stated in *Equation 6*, the equation for the velocity potential *ϕ*_*ρ*_ is

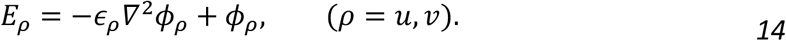

Again, 0 < *∈*_*ρ*_ < 1.

### The dimensionless model

Non-dimensionalisation, the removal of physical dimensions (units) of the variables, gives critical insight into the typical values and the relative magnitudes of the parameters that produce biologically reasonable solutions. Thus, we define the following non-dimensional variables:

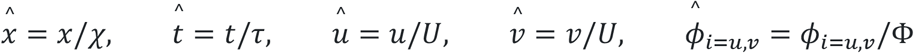

where *U* is a normalising density and Φ is a normalising velocity potential *ϕ*_*u,v*_. Setting Φτ/*χ*^2^=1 and *r*τ=1 and dropping primes give the non-dimensionalised system:

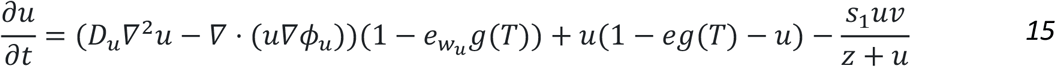

and

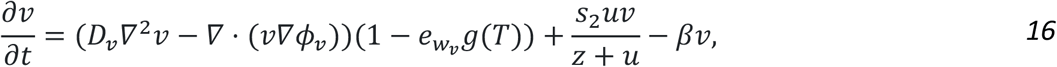

with the dimensionless equations for *E*_*u*_ and *E*_*v*_

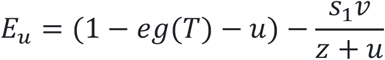

and

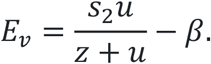

The non-dimensional equations for *ϕ*_*u*_ and *ϕ*_*v*_ are given by

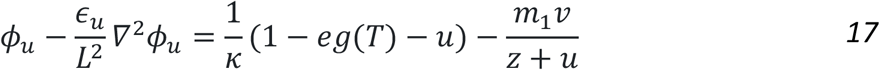

and

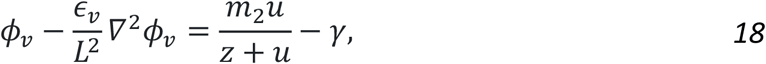

where

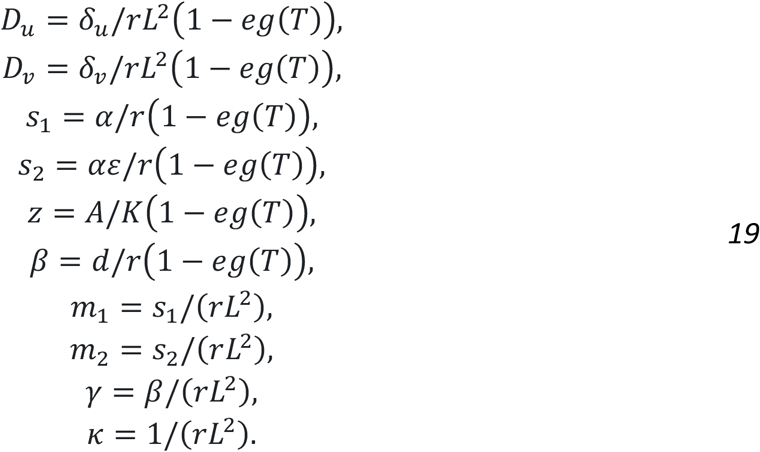

### Model stability analysis of the dimensionless model

We follow by carrying out a linear stability analysis of the temperature-dependent (*Equations 15 and 16*) to determine the equilibrium point of the spatially homogeneous model by switching off the spatial derivatives giving rise to the following movement-free system

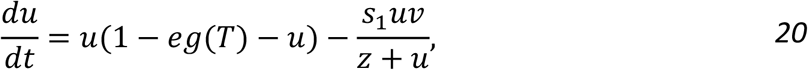

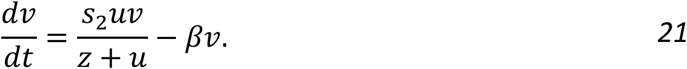

Setting the time derivative equal to zero, we observe the following three nonnegative equilibrium states for the temperature-dependent movement-free model:

- the unstable trivial state, (0,0), with eigenvalues 1 − *eg*(*T*) and −*β*. Since 0 ≤ *eg*(*T*) ≤ 1, (0,0) is an unstable saddle point.
- the locally asymptotically stable host-only state, (1 − *eg*(*T*),0), where both eigenvalues are negative if the condition

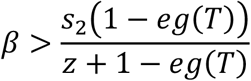 holds, or else (1 − *eg*(*T*),0) is an unstable saddle point.
- the coexistent state, (*u*^∗^, *v*^∗^), where

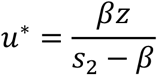

and

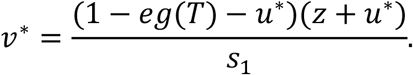

This steady state is only feasible if

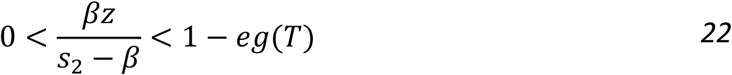

and the positive equilibrium point (*u*^∗^, *v*^∗^) is locally asymptotically stable if

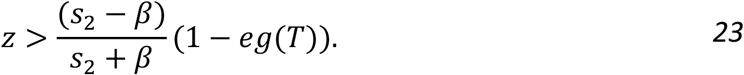

The condition in *Equation 22* gives the criterion expression for the feasibility of the coexistence steady state (*u*^∗^, *v*^∗^) (i.e., *u*^∗^ > 0 and *v*^∗^ > 0) which implies that the coexistence steady state is only feasible for a range of favourable temperatures and the size of this range depends on the values of *β, z, s*_*2*_ and *e*. Outside this range the coexistence steady state is not feasible anymore, giving rise to the host-only state (1 − *eg*(*T*), 0) which is stable for those very low or very high temperatures. Therefore, the climate function is set to change the dynamics of the predator-prey system. We have observed that the equilibrium point (0,0) is always unstable for positive parameters. Some kind of bifurcation may occur at (1 − *eg*(*T*), 0) but we examine the Hopf bifurcation near the coexisting steady state (*u*^∗^, *v*^∗^) to illustrate the outcomes which are beyond the theoretical finding (see Supplementary Material).

### Case Study: Dynamics of a parasitoid-butterfly system in the Iberian Peninsula and parameter values

We use the proposed temperature-dependent two-dimensional diffusion-advection-reaction model (*Equations 15-19*) described above to explore the effect of warmer summers and predicted increases in extreme temperatures (Alexander et al., 2006; Meehl & Tebaldi, 2004; Stocker & Qin, 2013) on the spatial dynamics of a parasitoid-butterfly system. Insects are well-suited to explore the stability and evolution of their population dynamics in the face of global warming. Like other ectotherms, insects cannot thermoregulate to buffer variation in temperature, which makes them particularly sensitive to climate change. Temperature shapes many aspects of insects’ biology, including their development and movement (e.g., Audusseau et al., 2013; Sheridan & Bickford, 2011), and generally, these responses follow a bell-shaped relationship (*Figure 1*b). For species involved in strong trophic interactions, like parasitoids that rely on their hosts to survive and reproduce, climate change can affect their ability to adapt to ongoing changes by altering the spatio-temporal determinants of the interaction. For insects, a change in temperature extreme is probably even more important as it has a direct impact on their survival (Ma et al., 2021).

We parameterise our model to forecast the dynamics of nettle-feeding butterflies and their parasitoids, a system that we have intensively monitored in the field (Audusseau et al. 2020). We explore their dynamics in the Iberian Peninsula under different past and future climatic conditions. The Iberian Peninsula offers a wide range of temperatures, including warm temperatures which are likely to be close, or beyond, the tolerance range of our species (the parasitoid and their host), especially as it also matches the hotter margin of nettle-feeding butterflies’ distribution (Settele et al., 2008). At the hot range margins, any even small changes may drastically alter species population dynamics (Evans et al., 2022). The choice of the model parameters (*Table 1*) was strongly based on Pearce et al. (2006) and checked for their consistency with our field observations for these species (Audusseau et al., 2017, 2020, 2021). Further, host dispersal is assumed to be greater than the dispersal of its parasitoid species. Indeed, parasitoid species are generally smaller and disperse over shorter distances than their hosts, making the impact of climate change on dispersal ability likely more pronounced for the parasitoid than the host and, hence, 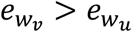.

**Table 1.** Parameters and values used for the numerical simulation of the temperature-dependent two-dimensional predator-prey model parasitoid-butterfly system in the Iberian Peninsula.

| Parameter | Value | Description |
| --- | --- | --- |
| $\delta_u$ | 0.01 | Host diffusive parameter |
| $\delta_v$ | 0.0003 | Parasitoid diffusive parameter |
| $r$ | 0.4 | Intrinsic growth rate for host |
| $K$ | 250 | Carrying capacity for host |
| $\varepsilon$ | 0.4 | Parasitoid conversion efficiencies of hosts to parasitoids |
| $\alpha$ | 0.35 | Parasitism rate |
| $d$ | 0.09 | Intrinsic mortality rate of parasitoids |
| $A$ | 80 | Parasitoid half-saturation constant |
| Constant | Value | Description |
| $e_{w_u}$ | 0.7 | Relative effect of temperature on host velocity |
| $e_{w_v}$ | 0.9 | Relative effect of temperature on parasitoid velocity |
| $e$ | 0.4 | Relative effect of temperature on host growth rate and carrying capacity |
| $\epsilon$ | 0.25 | smoothing constant |

We consider a spatial domain of the extent of the mainland portion of the Iberian Peninsula (Spain and Portugal), i.e., 1085 km East-West and 950 km North-South. Thus, the parameters in *Equation 19* for the dimensionless temperature-dependent model are given by

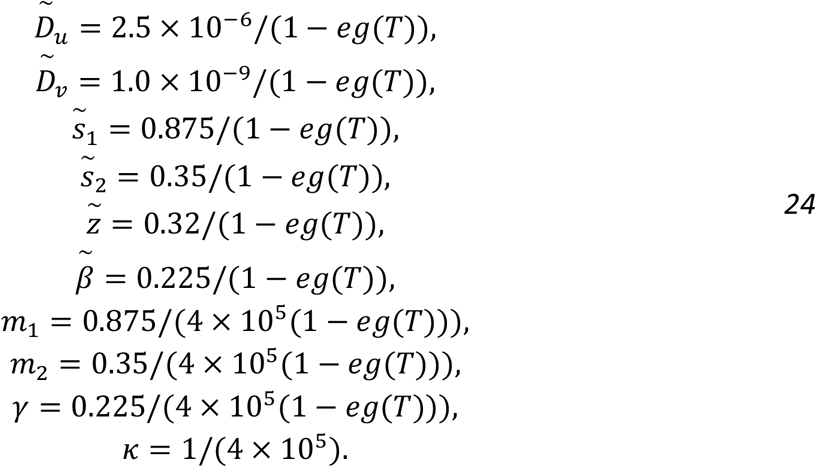

### Numerical solution

The dimensionless system (*Equations 15-18*) is approximated by the finite difference method and a forward Euler time-stepping method is used to integrate the equations (Supplementary Material). Second and first-order terms are approximated using central difference and one-sided approximation, respectively. For the solutions of *∂u/∂t* (*Equation 15*) and *∂v/∂t* (*Equation 16*), the value of the velocity potentials *ϕ*_*u,v*_ are required. Hence, *Equations 17* and *18* are approximated first which need the formation of pentadiagonal matrices. For the full application of the finite difference method to the system (*Equations 15-18*), see Supplementary Material.

The selected timestep value for the simulations is Δ*t* = 0.1. Following a convergence analysis, we observe a reduction in the error of the solution with the utilization of a finer mesh, leading the approximation generated by the explicit scheme to converge towards the exact solution. Given that the system of equations lacks an analytical solution, we employ a scenario with numerous spatial nodes as a reference for comparison against the approximate solution.

While augmenting mesh density results in a decay of approximation error and is pivotal for enhancing model accuracy, it introduces practical challenges. Specifically, the increased computational demand stemming from additional calculation steps mandated by the stability considerations yields a more computationally expensive scheme. The repeated experiments of the two-dimensional temperature-dependent model by adjusting the space step Δ*x* and time step Δ*t* ensure that further reductions in the space step had no significant effect on the numerical results.

Alongside the experiments for the stability and accuracy of the numerical method, we assess the ability of the model and the numerical method to produce consistent and reliable results across a range of different parameter sets and diverse initial conditions (Supplementary Material).

### Domain discretisation, boundaries and initial conditions

To apply the explicit finite difference scheme for approximating the system (*Equations 15-18*), we encoded the model on a rectangular domain defined by 0 ≤ *x* ≤ 108.5 and 0 ≤ *y* ≤ 95. We discretize the domain by 130×78 equally spaced nodes (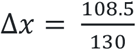 and 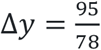), as shown in Figure 2. For the outer boundaries of the rectangular domain, we apply zero Neumann boundary conditions which confine species, while to create the boundaries separating land and sea (Figure 2a) we set all the parameters to zero at the sea, including the diffusion parameters.

The initial conditions used to model the response of our focal host-parasitoid system were the same for both species. The host and the parasitoid were set to initially occupy a region of the domain depicted in Figure 2b and defined by the function

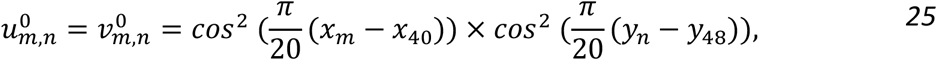

where *m* and *n* denote the intervals (*x*_30_, *x*_50_) and (*y*_38_, *y*_58_), respectively, while the initial conditions of *u* and *v*are zero outside these intervals.

### Climatic Data

We provide numerical approximations of the spatial dynamics of the parasitoid-butterfly system under four climatic environments, to explore the impact of past and future climates on these species. Historical and projected climate data were derived from high-resolution (0.11° × 0.11°) regional climate models (RCM) calculated over Europe (European domain) as part of the Coordinated Regional Climate Downscaling Experiment (CORDEX, https://cordex.org). We used historical data from 1951 and 2000 and projected data from the CORDEX climate projection experiment for the years 2020, 2050 and 2099 calculated under a future socio-economic climate scenario, namely RCP 4.5 (Copernicus Climate Change Service, Climate Data Store, 2019). To ensure consistency between the historical and future climate data, we used the simulations from the GCM: MOHC-HadGEM2-ES, RCM: CLMcom-CLM-CCLM4-8-17 and the ensemble: r1i1p1 (see documentation in Copernicus Climate Change Service, Climate Data Store, 2019). Using the same ensemble for the historical and projected climate models, the results of the output of historical models were used to form the basis for future climate projections. From the output of these climate experiments (historical and projected), we used the average of the maximum daily air temperature near the surface (2 m above the ground) observed between March and November. This metric provides the temperature maxima during the period when most butterfly species disperse and reproduce in southwest Europe and allows us to explore the impact of temperature extremes on the local and regional dynamics of butterfly parasitoid systems.

## Results

Using the initial conditions of *Equation 25* and the parameter values in *Equation 24*, we found that over the domain of the Iberian Peninsula, both populations reached stability at t ≈ 6000 (*Figure 3*). In all four scenarios, the butterfly population reached stability faster than their parasitoid counterpart (*Figure 3*). As the dynamics of parasitoid populations mathematically (and biologically) rely on the dynamics of their hosts, this delay mainly reflects the time needed for the parasitoid population to stabilize after its hosts. The outcomes obtained from the four distinct climatic landscapes showed species-specific population dynamics, resulting from the chosen parameter values (*Figure 3*). The results highlight the asymmetric impact of increasing temperatures on the population dynamics of these two species. The marginal increase in the total prey population is most likely explained by the approximate halving of the predator population between the years 1951, 2020, 2050 and 2099. This phenomenon arises from the intentional bias in temperature effects, favouring prey dispersal over predator dispersal. Consequently, prey can more effectively escape regions of elevated temperatures or high predator density.

**Figure 3.**
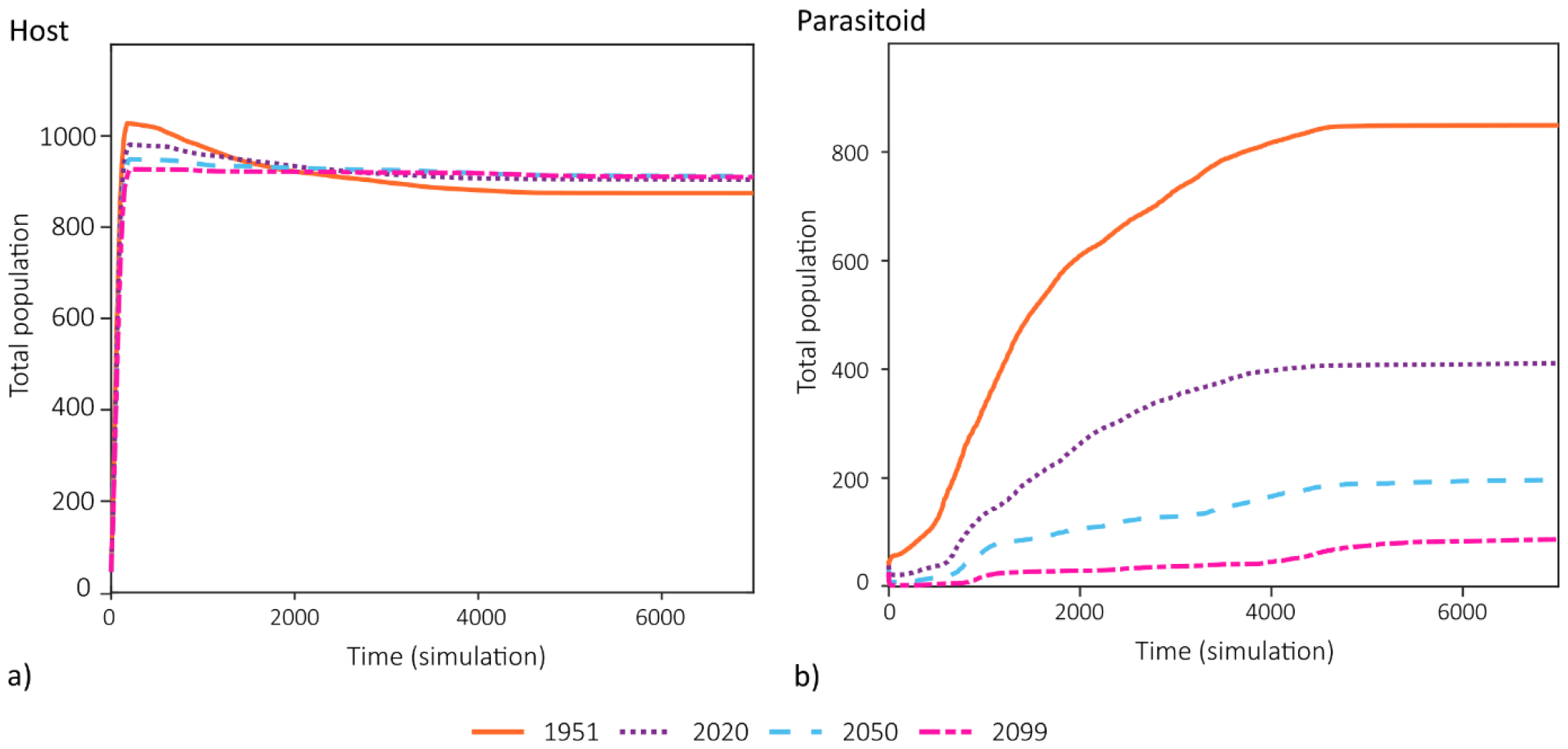
Temporal evolution of the total a) host population (u) and b) parasitoid population (v) throughout the simulations, according to four distinct climate data considered, being the average maximum daily temperature reported in years 1951, 2020, and predicted for 2050 and 2099.

Across the Iberian Peninsula, regions of high parasitoid densities are also regions with lower average daily maximum temperatures during the flying season for these species (Figure 4). The butterfly population is widespread throughout the domain and at relatively high densities. Yet, it reaches highest densities in areas where the parasitoid is absent and where the average daily maximum temperatures are lower. A comparative analysis across the years distinctly highlights a consistent trend of diminishing species expansion with increasing temperatures. Notably, for the host population we observe fewer areas with high population densities while clusters of parasitoids are predominantly found in the northern part of the domain, particularly in the results of the years 2050 and 2099. Remarkably, both species are stable globally (over the entire domain) and locally (over each mesh cell) after a certain number of time iterations. The observations align seamlessly with the results of the stability analysis revealing that the coexistence steady state remains stable within a favourable temperature range (Figure 4). Outside this range, the only stable steady state is characterized by the presence of the host population alone. This spatial pattern underscores the sensitivity of the studied ecological system to temperature extremes, highlighting the impact of temperature on species coexistence.

**Figure 4.**
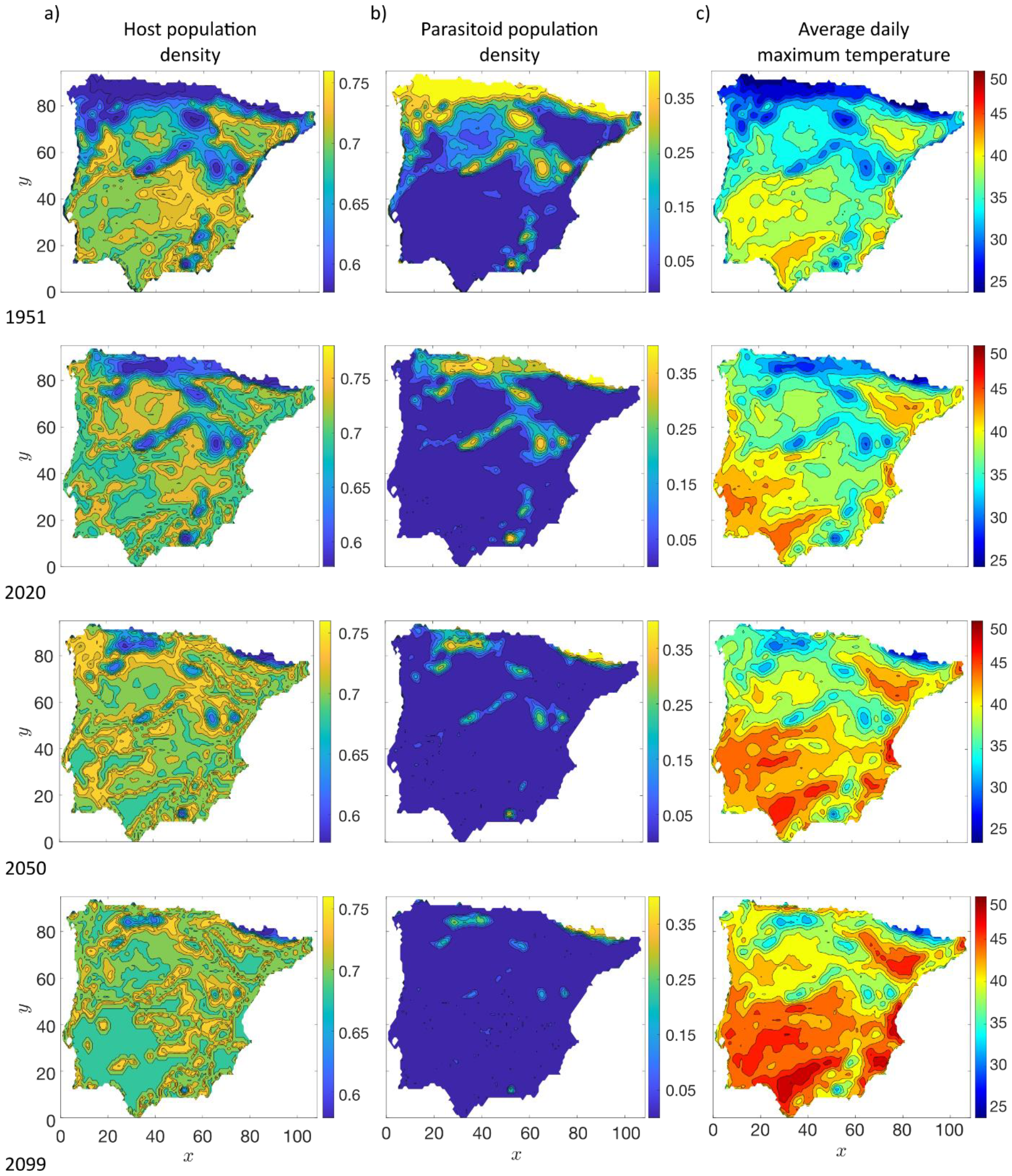
Spatial distributions of the population densities (ranging between 0 and 1) of a) the host butterfly and b) the parasitoid wasp at stability (t = 12000), modelled for the years 1951, 2020, 2050 and 2099. c) Historical (1951 and 2020) and predicted (2050 and 2099) average maximum daily temperatures between March and November, fitted for future socio-economic climate scenario RCP 4.5.

Different sets of parameters representing diverse ecological scenarios were incorporated into the model to simulate the population dynamics of predators and prey (Supplementary Material, Figure 3). The numerical approximations produced biologically relevant outcomes, illustrating consistent responses to alterations in parameters. Alongside testing with varied parameter values, we assessed the reliability of the model and numerical method through different initial conditions. We found that the model’s behaviour is reflective of the complexities inherent in real-world systems and the numerical method is capable of handling various parameter sets and initial conditions without instability. Our findings confirm the robustness of both the model and the numerical method.

## Discussion

We have developed a spatially explicit reaction-diffusion-advection model for predator-prey systems that integrates the influence of local conditions on species reproduction rate, local carrying capacity and individual dispersal. By explicitly accounting for species-specific responses to climate variables, our model enables its user to explore the dynamics of predator-prey interactions in changing environments.

The numerical simulations presented for our case study, using the finite difference method, demonstrate the use of our model to investigate and predict the distribution and dynamics of a parasitoid-host system in a two-dimensional spatial domain with realistic landscape under past, current and future climates. These simulations show dynamics and changes in species distribution that are consistent with our empirical understanding of the parasitoid-host systems associated with nettle-feeding butterflies and their parasitoids (Audusseau et al., 2017, 2020, 2021). The dynamics of parasitoids and hosts exhibit a stable oscillatory coexistence for a range of favourable temperatures. Above or below this temperature range, the stable state of coexistence becomes unstable, leading to parasitoid extinction and the spread of the hosts throughout the domain where thermal conditions are suitable. While an increase in average maximum temperature affects the distribution of both species, the effect of warming was more important for the parasitoids, as their abundance decreased more than 10-fold between 1951 and 2099. The asymmetric effect of climate change on host-parasitoid interactions is consistent with the results reported in other studies (Furlong & Zalucki, 2017; Godfray et al., 1994). However, reverse patterns have also been reported, in which an increase in temperature is more lethal for the prey than for their predators (Cavigliasso et al., 2021; Thomas & Blanford, 2003).

While the numerical method used in this paper has been used for various problems and has been proven to be highly robust and reliable, considerations of such methods, known as fixed mesh methods, arise regarding their accuracy, especially for problems where the solution exhibits a high degree of spatial activity and where regions of rapid variation of the solution move in space as time evolves. In such cases, having a mesh that moves with time would be desirable to track these features accurately without the computational expense of increasing the resolution everywhere. Applications of a moving mesh finite difference method on population dynamics for one-dimensional spatial domains can be found in Baines & Christou (2021, 2022), with ongoing work for two-dimensional domains.

The climate function that we have developed and integrated into a two-dimensional domain is a significant improvement over previous reaction-diffusion-advection models (Grindrod, 1991; Kurowski et al., 2017; Taylor et al., 2020) and addresses some important limitations of correlative approaches commonly used in ecology, e.g., species distribution models (Araújo et al., 2019). By accounting for species-specific response curves to environmental variables, our model provides a powerful way to study the dynamics of biotic interactions and explore alternative scenarios. In contrast with other predictive models based on correlations, process-based population models present two main advantages. First, they are built on a mechanistic understanding of how the individual of a species responds to the environment and its consequence in terms of fitness, which ultimately link to the dynamics and distribution of populations. Second, the performance of process-based models is less dependent on the amount and resolution of available data (Howard et al., 2014), which makes them well-suited for predicting systems in regions or environmental conditions where little empirical data is available. The local and regional stability of the numerical approximations highlights the capability of our model to produce interpretable multiscale predictions that can be used to identify and evaluate risks associated with potential threats and assess species’ population dynamics and vulnerability locally and across their range.

To conclude, the numerical stability and the ecological relevance of the predictions derived from the simulations show that our model and the numerical scheme can be applied to other species and environmental variables. With the specific aim of better predicting the impact of environmental changes on biodiversity, the model could be applied to explore long-term impact and transient dynamics resulting from ongoing change such as the modification in the composition and structure of the landscape and change in the duration, frequency and intensity of extreme events. Such environmental changes can shape dispersal and influence the rescue effect in regions where species densities are declining, resulting in cascading changes in the predator-prey system. By linking population dynamics, biotic interactions, dispersal and species-specific responses to environmental change, our approach provides crucial insights into biological systems to support decision-making.

## Acknowledgements

Special thanks go to Mike Baines and Peter Sweby for their useful comments and suggestions on this work. Christou is grateful for the support from the UK Natural Environment Research Council [grant number NE/P012345/1]. Audusseau acknowledges support from the Swedish Research Council (2016-06737) and the European Union’s Horizon 2020 research and innovation programme under the Marie Skłodowska-Curie grant agreement 899546 (ECOHEAT).

## Conflict of Interest Statement

The authors have no conflict of interest.

## Author Contributions

Christou, Schmucki and Audusseau conceived the ideas and designed the methodology; Christou, Schmucki and Audusseau collected the data; Christou developed the mathematical models and performed the simulations and analysed; Christou, Schmucki and Audusseau led and contributed equally to the writing of the manuscript. All authors gave final approval for publication.

## Notes

### Competing Interest Statement

The authors have declared no competing interest.

## References

Alexander, L. V., Zhang, X., Peterson, T. C., Caesar, J., Gleason, B., Klein Tank, A. M. G., Haylock, M., Collins, D., Trewin, B., Rahimzadeh, F., Tagipour, A., Rupa Kumar, K., Revadekar, J., Griffiths, G., Vincent, L., Stephenson, D. B., Burn, J., Aguilar, E., Brunet, M., … Vazquez-Aguirre, J. L. (2006). Global observed changes in daily climate extremes of temperature and precipitation. Journal of Geophysical Research: Atmospheres, 111(D5), 2005JD006290. 10.1029/2005JD006290

Araújo, M. B., Anderson, R. P., Márcia Barbosa, A., Beale, C. M., Dormann, C. F., Early, R., Garcia, R. A., Guisan, A., Maiorano, L., Naimi, B., O’Hara, R. B., Zimmermann, N. E., & Rahbek, C. (2019). Standards for distribution models in biodiversity assessments. Science Advances, 5(1), eaat4858. 10.1126/sciadv.aat4858

Audusseau, H., Baudrin, G., Shaw, M. R., Keehnen, N. L. P., Schmucki, R., & Dupont, L. (2020). Ecology and Genetic Structure of the Parasitoid Phobocampe confusa (Hymenoptera: Ichneumonidae) in Relation to Its Hosts, Aglais Species (Lepidoptera: Nymphalidae). Insects, 11(8), Article 8. 10.3390/insects11080478

Audusseau, H., Le Vaillant, M., Janz, N., Nylin, S., Karlsson, B., & Schmucki, R. (2017). Species range expansion constrains the ecological niches of resident butterflies. Journal of Biogeography, 44(1), 28–38. 10.1111/jbi.12787

Audusseau, H., Nylin, S., & Janz, N. (2013). Implications of a temperature increase for host plant range: Predictions for a butterfly. Ecology and Evolution, 3(9), 3021–3029. 10.1002/ece3.696

Audusseau, H., Ryrholm, N., Stefanescu, C., Tharel, S., Jansson, C., Champeaux, L., Shaw, M. R., Raper, C., Lewis, O. T., Janz, N., & Schmucki, R. (2021). Rewiring of interactions in a changing environment: Nettle-feeding butterflies and their parasitoids. Oikos, 130(4), 624–636. 10.1111/oik.07953

Baines, M. J., & Christou, K. (2021). A Moving-Mesh Finite-Difference Method for Segregated Two-Phase Competition-Diffusion. Mathematics, 9(4), 386. 10.3390/math9040386

Baines, M. J., & Christou, K. (2022). A Numerical Method for Multispecies Populations in a Moving Domain Using Combined Masses. Mathematics, 10(7), 1124. 10.3390/math10071124

Bellard, C., Bertelsmeier, C., Leadley, P., Thuiller, W., & Courchamp, F. (2012). Impacts of climate change on the future of biodiversity. Ecology Letters, 15(4), 365–377. 10.1111/j.1461-0248.2011.01736.x

Berg, M. P., Kiers, E. T., Driessen, G., Van Der Heijden, M., Kooi, B. W., Kuenen, F., Liefting, M., Verhoef, H. A., & Ellers, J. (2010). Adapt or disperse: Understanding species persistence in a changing world. Global Change Biology, 16(2), 587–598. 10.1111/j.1365-2486.2009.02014.x

Blois, J. L., Zarnetske, P. L., Fitzpatrick, M. C., & Finnegan, S. (2013). Climate Change and the Past, Present, and Future of Biotic Interactions. Science, 341(6145), 499–504. 10.1126/science.1237184

Boggs, C. L. (2016). The fingerprints of global climate change on insect populations. Current Opinion in Insect Science, 17, 69–73. 10.1016/j.cois.2016.07.004

Cantrell, R. S., Cosner, C., Lou, Y., & Xie, C. (2013). Random dispersal versus fitness-dependent dispersal. Journal of Differential Equations, 254(7), 2905–2941. 10.1016/j.jde.2013.01.012

Cavigliasso, F., Gatti, J.-L., Colinet, D., & Poirié, M. (2021). Impact of Temperature on the Immune Interaction between a Parasitoid Wasp and Drosophila Host Species. Insects, 12(7), 647. 10.3390/insects12070647

Copernicus Climate Change Service, Climate Data Store. (2019). CORDEX regional climate model data on single levels [NetCDF]. 10.24381/cds.bc91edc3

Cosner, C., & Winkler, M. (2014). Well-posedness and qualitative properties of a dynamical model for the ideal free distribution. Journal of Mathematical Biology, 69(6), 1343–1382. 10.1007/s00285-013-0733-z

Dawes, J. H. P., & Souza, M. O. (2013). A derivation of Holling’s type I, II and III functional responses in predator–prey systems. Journal of Theoretical Biology, 327, 11–22. 10.1016/j.jtbi.2013.02.017

DeLong, J. P., & Lyon, S. (2020). Temperature alters the shape of predator–prey cycles through effects on underlying mechanisms. PeerJ, 8, e9377. 10.7717/peerj.9377

Evans, L. C., Melero, Y., Schmucki, R., Boersch-Supan, P. H., Brotons, L., Fontaine, C., Jiguet, F., Kuussaari, M., Massimino, D., Robinson, R. A., Roy, D. B., Schweiger, O., Settele, J., Stefanescu, C., Turnhout C. A. M. van, & Oliver, T. H. (2022). Bioclimatic context of species’ populations determines community stability. Global Ecology and Biogeography. 10.1111/geb.13527

Furlong, M. J., & Zalucki, M. P. (2017). Climate change and biological control: The consequences of increasing temperatures on host–parasitoid interactions. Current Opinion in Insect Science, 20, 39–44. 10.1016/j.cois.2017.03.006

Fusco, G., & Minelli, A. (2010). Phenotypic plasticity in development and evolution: Facts and concepts. Philosophical Transactions of the Royal Society B: Biological Sciences, 365(1540), 547–556. 10.1098/rstb.2009.0267

Gentleman, W., Leising, A., Frost, B., Strom, S., & Murray, J. (2003). Functional responses for zooplankton feeding on multiple resources: A review of assumptions and biological dynamics. Deep Sea Research Part II: Topical Studies in Oceanography, 50(22), 2847–2875. 10.1016/j.dsr2.2003.07.001

Godfray, H. C. J., Hassell, M. P., & Holt, R. D. (1994). The Population Dynamic Consequences of Phenological Asynchrony between Parasitoids and their Hosts. Journal of Animal Ecology, 63(1), 1–10. 10.2307/5577

Gretchko, S., Marley, J., & Tyson, R. C. (2018). The Effects of Climate Change on Predator-Prey Dynamics. 10.48550/ARXIV.1805.11816

Grindrod, P. (1988). Models of individual aggregation or clustering in single and multi-species communities. Journal of Mathematical Biology, 26(6), 651–660. 10.1007/BF00276146

Grindrod, P. (1991). Patterns and waves: The theory and applications of reaction-diffusion equations. Clarendon Press ; Oxford University Press.

Hassell, M. P., Lawton, J. H., & Beddington, J. R. (1977). Sigmoid Functional Responses by Invertebrate Predators and Parasitoids. The Journal of Animal Ecology, 46(1), 249. 10.2307/3959

Holling, C. S. (1959). Some Characteristics of Simple Types of Predation and Parasitism. The Canadian Entomologist, 91(7), 385–398. 10.4039/Ent91385-7

Howard, C., Stephens, P. A., Pearce-Higgins, J. W., Gregory, R. D., & Willis, S. G. (2014). Improving species distribution models: The value of data on abundance. Methods in Ecology and Evolution, 5(6), 506–513. 10.1111/2041-210X.12184

Jeschke, J. M., Kopp, M., & Tollrian, R. (2002). Predator Functional Responses: Discriminating Between Handling and Digesting Prey. Ecological Monographs, 72(1), 95–112. 10.1890/0012-9615(2002)072[0095:PFRDBH]2.0.CO;2

Kurowski, L., Krause, A. L., Mizuguchi, H., Grindrod, P., & Van Gorder, R. A. (2017). Two-Species Migration and Clustering in Two-Dimensional Domains. Bulletin of Mathematical Biology, 79(10), 2302–2333. 10.1007/s11538-017-0331-0

Laws, A. N. (2017). Climate change effects on predator–prey interactions. Current Opinion in Insect Science, 23, 28–34. 10.1016/j.cois.2017.06.010

Lenoir, J., Bertrand, R., Comte, L., Bourgeaud, L., Hattab, T., Murienne, J., & Grenouillet, G. (2020). Species better track climate warming in the oceans than on land. Nature Ecology & Evolution, 4(8), Article 8. 10.1038/s41559-020-1198-2

Lotka, A. J. (1920). Analytical Note on Certain Rhythmic Relations in Organic Systems. Proceedings of the National Academy of Sciences, 6(7), 410–415. 10.1073/pnas.6.7.410

Lou, Y., & Ni, W.-M. (1996). Diffusion, Self-Diffusion and Cross-Diffusion. Journal of Differential Equations, 131(1), 79–131. 10.1006/jdeq.1996.0157

Ma, C.-S., Ma, G., & Pincebourde, S. (2021). Survive a Warming Climate: Insect Responses to Extreme High Temperatures. Annual Review of Entomology, 66(1), 163–184. 10.1146/annurev-ento-041520-074454

Meehl, G. A., & Tebaldi, C. (2004). More Intense, More Frequent, and Longer Lasting Heat Waves in the 21st Century. Science, 305(5686), 994–997. 10.1126/science.1098704

Parmesan, C. (2007). Influences of species, latitudes and methodologies on estimates of phenological response to global warming. Global Change Biology, 13(9), 1860–1872. 10.1111/j.1365-2486.2007.01404.x

Pearce, I. G., Chaplain, M. A. J., Schofield, P. G., Anderson, A. R. A., & Hubbard, S. F. (2006). Modelling the spatio-temporal dynamics of multi-species host–parasitoid interactions: Heterogeneous patterns and ecological implications. Journal of Theoretical Biology, 241(4), 876–886. 10.1016/j.jtbi.2006.01.026

Poloczanska, E. S., Brown, C. J., Sydeman, W. J., Kiessling, W., Schoeman, D. S., Moore, P. J., Brander, K., Bruno, J. F., Buckley, L. B., Burrows, M. T., Duarte, C. M., Halpern, B. S., Holding, J., Kappel, C. V., O’Connor, M. I., Pandolfi, J. M., Parmesan, C., Schwing, F., Thompson, S. A., & Richardson, A. J. (2013). Global imprint of climate change on marine life. Nature Climate Change, 3(10), Article 10. 10.1038/nclimate1958

Pounds, A. J., Bustamante, M. R., Coloma, L. A., Consuegra, J. A., Fogden, M. P. L., Foster, P. N., La Marca, E., Masters, K. L., Merino-Viteri, A., Puschendorf, R., Ron, S. R., Sánchez-Azofeifa, G. A., Still, C. J., & Young, B. E. (2006). Widespread amphibian extinctions from epidemic disease driven by global warming. Nature, 439(7073), Article 7073. 10.1038/nature04246

Rall, B. C., Vucic-Pestic, O., Ehnes, R. B., Emmerson, M., & Brose, U. (2010). Temperature, predator– prey interaction strength and population stability. Global Change Biology, 16(8), 2145–2157. 10.1111/j.1365-2486.2009.02124.x

Rowell, J. T. (2009). The Limitation of Species Range: A Consequence of Searching Along Resource Gradients. Theoretical Population Biology, 75(2–3), 216–227. 10.1016/j.tpb.2009.03.001

Segel, L. A., & Jackson, J. L. (1972). Dissipative structure: An explanation and an ecological example. Journal of Theoretical Biology, 37(3), 545–559. 10.1016/0022-5193(72)90090-2

Sekerci, Y. (2020). Climate change effects on fractional order prey-predator model. Chaos, Solitons & Fractals, 134, 109690. 10.1016/j.chaos.2020.109690

Sekerci, Y., & Petrovskii, S. (2015). Mathematical Modelling of Plankton–Oxygen Dynamics Under the Climate Change. Bulletin of Mathematical Biology, 77(12), 2325–2353. 10.1007/s11538-015-0126-0

Settele, J., Kudrna, O., Harpke, A., Kühn, I., Van Swaay, C., Verovnik, R., Warren, M., Wiemers, M., Hanspach, J., Hickler, T., Kühn, E., Van Halder, I., Veling, K., Vliegenthart, A., Wynhoff, I., & Schweiger, O. (2008). Climatic Risk Atlas of European Butterflies. BioRisk, 1, 1–712. 10.3897/biorisk.1

Sheridan, J. A., & Bickford, D. (2011). Shrinking body size as an ecological response to climate change. Nature Climate Change, 1(8), Article 8. 10.1038/nclimate1259

Shigesada, N., Kawasaki, K., & Teramoto, E. (1979). Spatial segregation of interacting species. Journal of Theoretical Biology, 79(1), 83–99. 10.1016/0022-5193(79)90258-3

Skalski, G. T., & Gilliam, J. F. (2001). Functional Responses with Predator Interference: Viable Alternatives to the Holling Type II Model. Ecology, 82(11), 3083–3092. 10.2307/2679836

Stocker, T., & Qin, D. (Eds.). (2013). Climate change 2013: The physical science basis: summary for policymakers, a report of working group I of the IPCC: technical summary, a report accepted by working group I of the IPCC but not approved in detail: and frequently asked questions: part of the working group I contribution to the fifth assessment report of the intergovernmental panel on climate change. WMO, UNEP.

Taylor, N. P., Kim, H., Krause, A. L., & Van--Gorder, R. A. (2020). A Non-local Cross-Diffusion Model of Population Dynamics I: Emergent Spatial and Spatiotemporal Patterns. Bulletin of Mathematical Biology, 82(8), 112. 10.1007/s11538-020-00786-z

Thomas, M. B., & Blanford, S. (2003). Thermal biology in insect-parasite interactions. Trends in Ecology & Evolution, 18(7), 344–350. 10.1016/S0169-5347(03)00069-7

Traill, L. W., Lim, M. L. M., Sodhi, N. S., & Bradshaw, C. J. A. (2010). Mechanisms driving change: Altered species interactions and ecosystem function through global warming. Journal of Animal Ecology, 79(5), 937–947. 10.1111/j.1365-2656.2010.01695.x

Turing. (1952). The chemical basis of morphogenesis. Philosophical Transactions of the Royal Society of London. Series B, Biological Sciences, 237(641), 37–72. 10.1098/rstb.1952.0012

Visser, M. E., & Holleman, L. J. M. (2001). Warmer springs disrupt the synchrony of oak and winter moth phenology. Proceedings of the Royal Society of London. Series B: Biological Sciences, 268(1464), 289–294. 10.1098/rspb.2000.1363

Volterra, V. (1926). Fluctuations in the Abundance of a Species considered Mathematically1. Nature, 118(2972), 558–560. 10.1038/118558a0

Watkinson, A. R., & Gill, J. A. (2002). Climate change and dispersal. In British Ecological Society, J. M. Bullock, R. Kenward, & R. Hails (Eds.), Dispersal ecology: The 42nd symposium of the British Ecological Society held at the University of Reading, 2-5 April 2001 (1st ed, pp. 45–56). Blackwell Pub.

Wisz, M. S., Pottier, J., Kissling, W. D., Pellissier, L., Lenoir, J., Damgaard, C. F., Dormann, C. F., Forchhammer, M. C., Grytnes, J.-A., Guisan, A., Heikkinen, R. K., Høye, T. T., Kühn, I., Luoto, M., Maiorano, L., Nilsson, M.-C., Normand, S., Öckinger, E., Schmidt, N. M., … Svenning, J.-C. (2013). The role of biotic interactions in shaping distributions and realised assemblages of species: Implications for species distribution modelling. Biological Reviews, 88(1), 15–30. 10.1111/j.1469-185X.2012.00235.x

