## Supplementary Material for "Modelling the spatiotemporal dynamics of multispecies population interactions in the context of climate change"

### Modelling the spatiotemporal dynamics of multi-species population interactions in the context of climate change : Supplementary Material

#### 0.1 Numerical simulations of the ODE system

From the model analysis of the movement-free temperature-dependent ODE system for the predator-prey model with Holling-type II functional response (Equations 20 and 21), we have gained insights into the parameter values required to produce biologically realistic behaviour by looking at the three non-negative equilibrium steady states. Here, special attention is given to the coexistence state $(u^*, v^*)$ where by choosing parameter $z$ as a bifurcation parameter, sufficient conditions for the existence of Hopf bifurcation have been presented using numerical simulations. The conditions derived in this section regarding the range of parameter values which must be used to achieve a stable coexistence steady state will be used when we numerically approximate the reaction-diffusion-advection system (Equations 15-19).

We numerically approximate Equations 20 and 21 with initial conditions $u(0) = 0.7$ and $v(0) = 0.5$. We fix the following parameters:

$$\beta = 0.175, \quad s_1 = 0.875, \quad s_2 = 0.35, \quad e = 0.25 \quad \text{and} \quad T = 27$$

while we take $z$ as the control parameter. The values of $\beta$, $s_1$ and $s_2$ are chosen based on the ones used by Pearce et al. (2006) where $r = 0.4$, $\alpha = 0.35$, $\varepsilon = 0.4$ and $d = 0.07$ while $T = 27$ corresponds to the most favourable temperature for the species. For this set of parametric values, the coexistence equilibrium point exists whenever $0 < z < 1$ (see Equation 22) and is locally asymptotically stable for $z > 0.33$ (see Equation 23). Let us define the critical value $\overline{z} = 0.33$. We intend to show that equilibrium point $(u^*, v^*)$ undergoes a Hopf bifurcation when the parameter $z$ crosses its critical value $\overline{z} = 0.33$.
We present Figures 1 and 2 as evidence of Hopf bifurcation at $z = \overline{z}$. It is observed by Figure 1 that the system of Equations 20 and 21 has a stable positive equilibrium point $(u^*, v^*) = (0.3751, 0.5359)$ for $z = 0.375$. Figure 1a represents the positive equilibrium that is locally asymptotically stable and 1b represents the phase-portrait of $(u^*, v^*) = (0.3751, 0.5359)$ that is locally asymptotically stable.
In Figure 2, we show that the coexisting equilibrium becomes unstable for some $z = 0.3 < \overline{z}$. We observe that solutions of $u(t)$ and $v(t)$ oscillate around $(u^*, v^*)$. This represents the occurrence of a limit cycle and the destabilisation of coexisting equilibria. Thus, Hopf bifurcation is verified for $z < \overline{z}$. Figure 2a

represents the stable and periodic oscillations around the positive equilibrium $(u^*, v^*) = (0.3751, 0.5359)$ and the phase plane diagram shown in Figure 2b represents the stable limit cycle around $(u^*, v^*)$ when $z = 0.3 < \overline{z} = 0.33$. The stability of the limit cycle is determined by the direction of the neighbouring trajectories. If the neighbouring trajectories are approaching the limit cycle, as time increases, then the limit cycle is stable. Otherwise, the limit cycle is unstable if the trajectories tend away from the limit cycle.
Hence from Figures 1 and 2 we observe that as we decrease the value of the control parameter $z$, the coexistence of prey-predator changes from stable equilibrium to stable oscillatory coexistence.

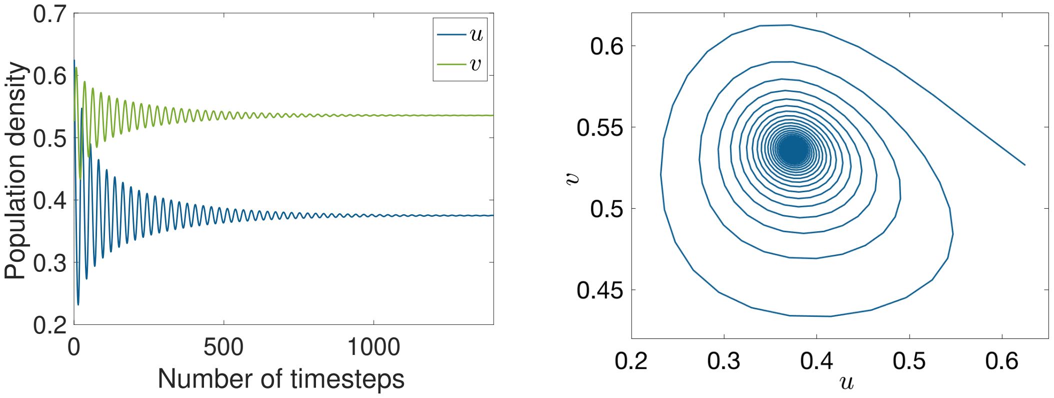


(a) Time evolution. (b) Phase-portait.

Figure 1: Stable coexisting equilibrium $(u^*, v^*) = (0.3751, 0.5359)$ for $z = 0.375 > \overline{z}$ for the system Equations 20 and 21.

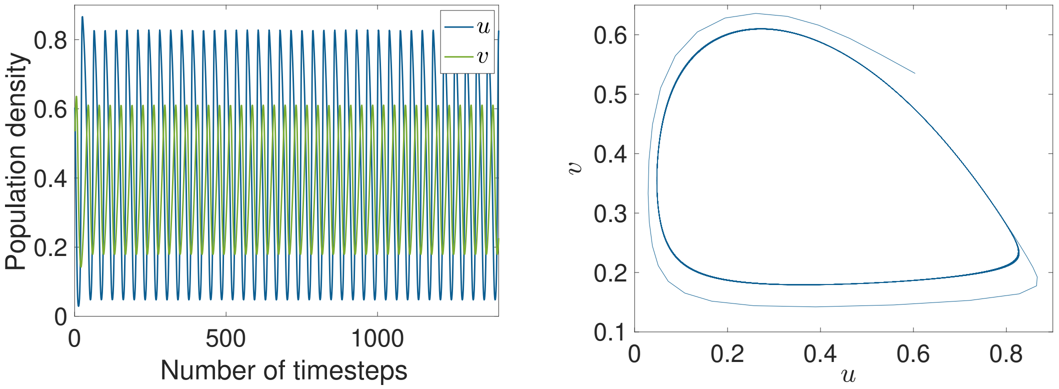


(a) Time evolution

(b) Phase-portait.

Figure 2: Stable limit cycle surrounding the coexisting equilibrium $(u^*, v^*) = (0.3751, 0.5359)$ for $z = 0.3 < \overline{z}$ for the system Equations 20 and 21.

#### 0.2 Finite difference scheme

The finite difference method stands out as a favored approach for tackling reaction-diffusion-advection equations, owing to its straightforward implementation, heightened efficiency, and accuracy. Extensive literature has already delved into the intricacies of such algorithms.

By discretizing both the spatial and temporal domains, the finite difference method transforms the challenge of solving for a continuous solution into a series of algebraic systems with finite dimensions. Explicit and implicit finite difference methods are generally applicable to solving time-dependent partial differential equations.

Due to the nonlinearity of the reaction terms in the equations of our model, a fully implicit scheme is inefficient for the approximation of the system 15-19. A semi-implicit scheme is possible for the solution of the system and can be derived by combining the two Equations 15 and 16 and constructing a sparse matrix equation which is solved to update the values of $u$ and $v$ at forward time levels. The operations of sparse matrices using standard dense-matrix structures and algorithms are slow and inefficient when applied to large sparse matrices as processing and memory are wasted on the zeros.

In this study, we employ the explicit finite difference method which is efficient in each temporal step compared to the implicit or semi-implicit ones but suffers stability issues. The stability consideration associated with the explicit scheme depends on the spatial and temporal discretisations. While a finer mesh enhances accuracy, it necessitates a smaller timestep to avert instability, implying the need for numerous time steps to track the solution across a sufficiently large time interval.

#### 0.3 Numerical solution

The dimensionless system Equations 15-19 is solved by the finite difference method and a forward Euler time-stepping method is used to integrate the equations. Second- and first-order terms are approximated using central difference and one-sided approximation, respectively. For the solutions of $\partial u/\partial t$ 15 and $\partial v/\partial t$ 16, the value of the velocity potentials $\phi_{u,v}$ are required. Hence, Equations 17 and 18 are approximated first which need the formation of pentadiagonal matrices. The equation approximating $\phi_u$ 17 after applying the finite difference approximations is

$$-\frac{\epsilon_u}{\Delta x^2}\phi^n_{u(i+1,j)} - \frac{\epsilon_u}{\Delta y^2}\phi^n_{u(i,j+1)} + \left(1 + \frac{2\epsilon_u}{\Delta x^2} + \frac{2\epsilon_u}{\Delta y^2}\right)\phi^n_{u(i,j)} -$$
$$\frac{\epsilon_u}{\Delta x^2}\phi^n_{u(i-1,j)} - \frac{\epsilon_u}{\Delta y^2}\phi^n_{u(i,j-1)} = \frac{1}{\kappa}\left(1 - eg(T_{(i,j)}) - u_{(i,j)}\right) - \frac{m_1 v^n_{(i,j)}}{z + u^n_{(i,j)}}$$
$$\text{for} \quad i = 2, ..., N \quad \text{and} \quad j = 2, ..., M$$

and for $\phi_v$ by Equation 18,

$$-\frac{\epsilon_v}{\Delta x^2}\phi^n_{v(i+1,j)} - \frac{\epsilon_v}{\Delta y^2}\phi^n_{v(i,j+1)} + \left(1 + \frac{2\epsilon_v}{\Delta x^2} + \frac{2\epsilon_v}{\Delta y^2}\right)\phi^n_{v(i,j)} -$$
$$\frac{\epsilon_v}{\Delta x^2}\phi^n_{v(i-1,j)} - \frac{\epsilon_v}{\Delta y^2}\phi^n_{v(i,j-1)} = \frac{m_2 u^n_{(i,j)}}{z + u^n_{(i,j)}} - \gamma$$
$$\text{for} \quad i = 2, ..., N \quad \text{and} \quad j = 2, ..., M.$$

The dimensionless Equation 15 for $\partial u/\partial t$ after the finite difference approximations takes the form

$$\frac{\partial u}{\partial t} =$$

$$\Bigg[-\frac{1}{\Delta x^2}\left(\frac{\left(u^n_{(i+1,j)}+u^n_{(i,j)}\right)\left(\phi^n_{u(i+1,j)}-\phi^n_{u(i,j)}\right)}{2}-\frac{\left(u^n_{(i,j)}+u^n_{(i-1,j)}\right)\left(\phi^n_{u(i,j)}-\phi^n_{u(i-1,j)}\right)}{2}\right)-$$

$$\frac{1}{\Delta y^2}\left(\frac{\left(u^n_{(i,j+1)}+u^n_{(i,j)}\right)\left(\phi^n_{u(i,j+1)}-\phi^n_{u(i,j)}\right)}{2}-\frac{\left(u^n_{(i,j)}+u^n_{(i,j-1)}\right)\left(\phi^n_{u(i,j)}-\phi^n_{u(i,j-1)}\right)}{2}\right)+$$

$$\delta_u\left(\frac{u^n_{(i-1,j)}-2u^n_{(i,j)}+u^n_{(i+1,j)}}{\Delta x^2}\right)+\delta_u\left(\frac{u^n_{(i,j-1)}-2u^n_{(i,j)}+u^n_{(i,j+1)}}{\Delta y^2}\right)\Bigg]\left(1-e_{w_u}g(T_{(i,j)})\right)+$$

$$u^n_{(i,j)}\left(1-eg(T_{(i,j)})-u^n_{(i,j)}\right)-\frac{s_1\,u^n_{(i,j)}\,v^n_{(i,j)}}{z+u^n_{(i,j)}}\qquad \text{for}\quad i=2,...,N\quad\text{and}\quad j=2,...,M.$$

and for $\partial v/\partial t$ by Equation 16 we get

$$\frac{\partial v}{\partial t} =$$

$$\Bigg[-\frac{1}{\Delta x^2}\left(\frac{\left(v^n_{(i+1,j)}+v^n_{(i,j)}\right)\left(\phi^n_{v(i+1,j)}-\phi^n_{v(i,j)}\right)}{2}-\frac{\left(v^n_{(i,j)}+v^n_{(i-1,j)}\right)\left(\phi^n_{v(i,j)}-\phi^n_{v(i-1,j)}\right)}{2}\right)-$$

$$\frac{1}{\Delta y^2}\left(\frac{\left(v^n_{(i,j+1)}+v^n_{(i,j)}\right)\left(\phi^n_{v(i,j+1)}-\phi^n_{v(i,j)}\right)}{2}-\frac{\left(v^n_{(i,j)}+v^n_{(i,j-1)}\right)\left(\phi^n_{v(i,j)}-\phi^n_{v(i,j-1)}\right)}{2}\right)+$$

$$\delta_v\left(\frac{v^n_{(i-1,j)}-2v^n_{(i,j)}+v^n_{(i+1,j)}}{\Delta x^2}\right)+\delta_v\left(\frac{v^n_{(i,j-1)}-2v^n_{(i,j)}+v^n_{(i,j+1)}}{\Delta y^2}\right)\Bigg]\left(1-e_{w_v}g(T_{(i,j)})\right)+$$

$$\frac{s_2\,u^n_{(i,j)}\,v^n_{(i,j)}}{z+u^n_{(i,j)}}-\beta v^n_{(i,j)}\qquad \text{for}\quad i=2,...,N\quad\text{and}\quad j=2,...,M.$$

The Equations for $\phi_u$, $\phi_v$, $\partial u/\partial t$ and $\partial u/\partial t$ at the outer boundaries are evaluated by Equations 17, 18, 15 and 16, respectively by accounting for zero Neumann boundary conditions.

For example, to evaluate $\phi_{u(1,1:M)}$ from Equation 17 (at the left-hand-side boundary, $x_1$, of Figure 2 or the main manuscript) we consider the zero Neumann boundary condition

$$\frac{\partial \phi_u}{\partial x}=0\quad \text{at}\quad x=0.$$

Applying a two-interval approximation gives

$$\frac{\phi_{u(2,1:M)} - \phi_{u(0,1:M)}}{2\Delta x} = 0$$

and hence

$$\phi_{u(0,1:M+1)} = \phi_{u(2,1:M+1)}.$$

#### 0.4 Fixed and Moving Meshes for Population Dynamics

The explicit finite difference numerical scheme used to approximate the parasitoid-host system is a standard numerical technique used by a lot of modellers to approximate a range of different PDEs. Even though the method is widely used and is easy to implement there are some drawbacks. We now briefly describe the limitations associated with accuracy considerations of the explicit finite difference method.

The leading term in the truncation error of a fixed-mesh explicit finite difference numerical scheme for a diffusion problem is proportional to $h\frac{d^4u}{dx^4}$, where $h = \frac{\delta \Delta x^2}{12}$, which is a numerical diffusion distinct from the diffusion intrinsic to the problem.

In convergence, as $\Delta x \to 0$ and hence $h$ tends to zero this numerical diffusion term also tends to zero (as it should) and the truncation error decreases as the leading term is eliminated. But if $\frac{d^4u}{dx^4}$ is large the magnitude is always large even when $h$ is small, and the results are adversely affected.

This situation occurs at a 'corner' of a near-discontinuity in $u$, for example at the base or top of a near-vertical part of a graph of $u$, particularly if it is moving.

In these circumstances, a superior description is to have a moving point at the discontinuity and replace the PDE by a jump condition, forming the so-called moving boundary. The moving mesh method based on conservation is well-suited to accommodate this description. The main feature of the moving mesh based on conservation is that moving nodes are concentrated in regions with increasing density $u$ since the method is based on a conservation principle that "forces" the mass (number of individuals of a population), i.e., the area under the graph of the density $u$, within each element of the discretised domain to remain constant for all time. Therefore, regions with increasing density $u$ will have a denser mesh as the area under the graph of $u$ within each element remains constant. Hence the derivative of the leading term in the truncation error balances out between the denominator and the numerator.

#### 0.5 Different dispersal climate factor

We carried out simulations for different sets of parameters representing diverse ecological scenarios. We investigate different parameter sets by varying one parameter each time, as shown in Figure 3 where we have set the climate factor for the host dispersal to be greater than that of its parasitoid while keeping the rest of the parameters constant. The results are found to be biologically relevant and consistent with respect to the parameter being varied in each experiment.

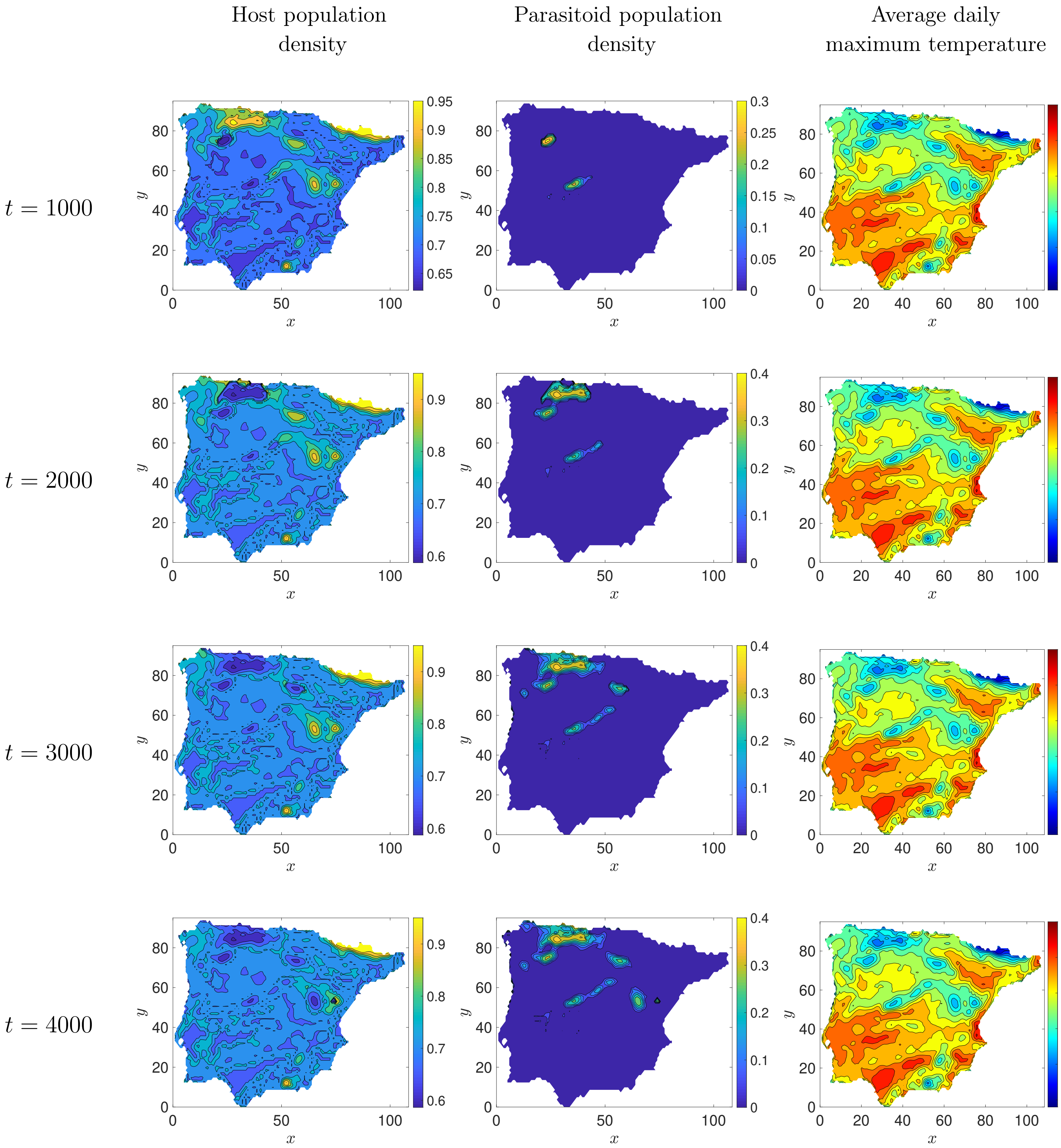
Host population density
Parasitoid population density
Average daily maximum temperature
t = 1000
t = 2000
t = 3000
t = 4000
x
y
0.95
0.9
0.85
0.8
0.75
0.7
0.65
0.6
0.3
0.25
0.2
0.15
0.1
0.05
0
0.4
50
45
40
35
30
25

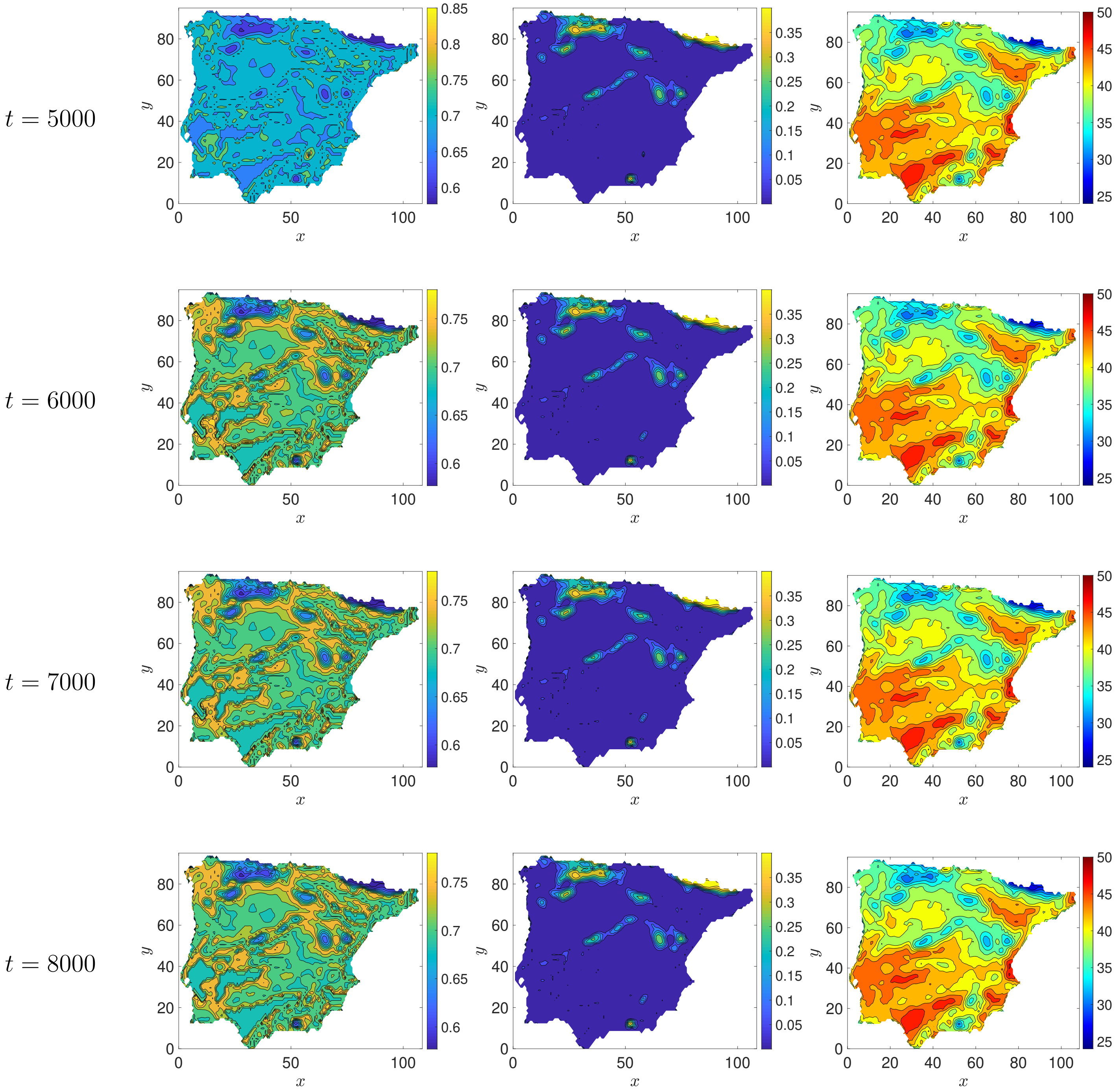


Figure 3: Simulations using the parameters in Equation 24 of the main manuscript but with different values of climate factor for the species dispersal. For this set of simulations, we set $e_{w_u} > e_{w_v}$ ($e_{w_u} = 0.9$ and $e_{w_v} = 0.7$).
